# Python Encoders for Archetypal Convex Hulls (PEACH): PyTorch-Based Archetypal Analysis

**DOI:** 10.64898/2025.12.29.696912

**Authors:** Alexander T. Honkala, Sanjay V. Malhotra

## Abstract

Samples of closely related cells often contain substantial cell state heterogeneity, which traditional clustering-based analyses struggle to de-convolve. Archetype analysis is an alternative analysis approach that identifies a minimal convex hull enclosing all data points in a dimensionally-reduced space, such as PCA space. The points of this convex manifold, or hull, each represent theoretical extremal states specialized in a subset of functions that are mutually exclusive to the extremal states found at each other vertex. The transition between two points represents a Pareto front of tradeoffs between potential cell states. Cell distances from each archetype and direction between archetypes then carry interpretable biological information revealing features of resource tradeoffs in cell state regulation. The archetypal points themselves are theoretical pure extremal states rarely accessed by real cells, which instead display a combination of archetype weights/mixture coefficients representing a mixture of specializations. By characterizing the relationships between gene expression and archetype distance, archetypal analysis can identify the phenotypic features of theoretically pure specialist states, the phenotypic tradeoffs of real cells, and the boundaries of accessible cell states in the analyzed sample. Here we present Python Encoders for Archetypal Convex Hulls (PEACH), a new PyTorch-based archetype analysis package compatible with the scVerse ecosystem that recapitulates and extends features available in previous archetype analysis packages. This includes a hyperparameter search function to identify the best *k* archetypes that fit an input dataset. PEACH uses a deliberately constrained autoencoder architecture to directly learn an archetypal latent space, replacing previous alternating least squares methods with a highly performant, GPU-accelerated archetype analysis method that is scalable to modern scRNAseq datasets of hundreds of thousands of cells. Crucially, PEACH includes automated hyperparameter search with cross-validation to identify optimal archetype and initialization configurations, enabling users to implement archetype analysis without extensive manual hyperparameter optimization and testing.

## INTRODUCTION

The adoption of statistical analysis techniques in bioscience has been constrained by computational feasibility, which drove the widespread use of techniques such as Principal Component Analysis (PCA), Uniform Manifold Approximation Projection (UMAP), and k-means clustering^1^. In contrast to clustering-based approaches that assign every cell to a group, archetype analysis instead identifies a globally convex hull enclosing the data in PCA space where hull features encode biologically interpretable features in terms of distance from each archetype and the archetype weight mixtures for each cell or datapoint characterized^2^. This inversion of the traditional clustering approaches enables users to identify what extremal states form boundaries on cell states accessible within the characterized sample, where the trajectories between archetypal extrema represent Pareto fronts of resource tradeoffs between mutually exclusive subsets of cell function. Distance from each archetype is then directly related to degree of specialization, with cells close to an archetype exhibiting a restricted, specialized state more associated with that archetype’s phenotypic characteristics than cells further away or more mixed between multiple archetypes^3^. In scRNAseq data with gene set activity scores, this can be used to identify both genes and pathways that drive archetype specialization, with mutually exclusive modules segregating cells into specialist phenotypes that reflect both the internal constraints and external pressures present at the time the cells were sampled^4^. These same principles can be applied to many other data types, including chemical structures, image analysis, and mass cytometry^5–10^. Importantly, when fit across multiple conditions pooled together, archetype analysis can reveal diverging trajectories within a convex hull for different conditions, where cells that all start at the same archetype mixture before perturbation follow distinct specialization paths in archetype space to different mixtures of specialist states. This reflects that cell state choices are not open-ended and are instead constrained both by prior state and tradeoffs in competing functions.

Archetypal analysis was originally formulated in 1994 by Breiman and Cutler^2^ as an alternating least squares method, requiring that each update solves several quadratic optimization terms that scale with dataset size as an *O*(*n*^2^) process^11^. This limitation has previously restricted the use of archetypal analysis to smaller datasets while more efficient techniques, such as non-negative matrix factorization with an (*O*(*m × n*)) scaling curve, were more broadly applied to emerging large biomedical datasets^1^. The development of ParTI in MATLAB by the Uri Alon group applied archetypal analysis to RNA sequencing data for the first time^3,4^, showing that it identifies extremal states that generalize across cancer types while exploring the application of archetypal analysis to morphological and evolutionary state spaces^5,6^. Other groups have applied it to image analysis, digital signal processing, and machine learning^8,12,13^. The following ParetoTI R package was more broadly used in finding cell differentiation-associated transcription factors in lung cancer^14^, intestinal crypt stem cell regulation^15^, liver zonation^3^, and transplant medicine^16–18^, where application of archetype analysis to histology images is under active development for the early identification of antibody-mediated transplant rejection^18,19^. Recent experiments with archetypal analysis, such as AAnet and MIDAA^20,21^, extended the archetype analysis approach to single-cell RNAseq and MNIST datasets. However, these approaches lack methods for hyperparameter search in identifying the *k* archetypes that best fit the data being characterized. Here, with Python Encoders for Archetypal Convex Hulls (PEACH), we introduce a new PyTorch-optimized package compatible with the broader scVerse ecosystem that recapitulates all features of ParTI and ParetoTI and is extensible to large modern scRNAseq datasets via GPU acceleration. PEACH achieves archetypal reconstruction *R*^2^ > 0.89 on benchmark datasets, demonstrating that archetypal mixing captures genuine biological state structure. Taking inspiration from several previous lines of archetype analysis research, PEACH uses a principal convex hull based initialization of archetype positions in a constrained variational autoencoder architecture in which the dimensions of the latent space are the learned archetypes. This creates a latent space that can be expressed in barycentric coordinates to enforce a geometric simplex structure, forming the archetypal convex hull, where every encoded cell is represented by a convex combination of archetypal weights/barycentric coordinates. Implementation of a Frobenius norm-based archetypal reconstruction measurement in the autoencoder loss term measures the sum of squared reconstruction errors across all cells and features to create smooth, differentiable gradients for effective training. Application of PEACH to a large scRNAseq dataset for circulating hematopoietic cells (HSCs)^22^ shows efficient exploration of multiple hyperparameter configurations, available as a systematic grid search, and reproducible training of its archetypal autoencoder model on datasets of varying sizes. Characterization of archetypal features recapitulates cell type clustering in a highly mixed subsample and, in a single cell type subsample, reveals novel phenotypes of plasmacytoid dendritic cells (pDCs). PEACH includes several features for phenotypic characterization of archetypes, including statistical testing for associated gene sets with detection of mutually exclusive patterns, a hypergeometric test for the association of categorical variables with archetype specialization, and a compatibility module for working with CellRank for trajectory inference.

### METHODS Model Architecture Overview

PEACH implements a VAE-based autoencoder, Deep_AA, with archetypal constraints on the learning process and latent space. The key constraint is that the latent space **z** directly represents archetypal coordinates (A matrix) for each cell, eliminating the need for a separate archetypal decomposition step. Taking as input PCA-reduced scRNAseq data, the input data matrix is formed *X* ∈ ℝ^*N×D*^ for N samples and D features (PCA dimensions), the archetype positions are represented by *Y* ∈ ℝ^*K×D*^ for K archetypes, archetype coordinates per cell are formed by *A* ∈ ℝ^*N×K*^, and the latent representation is set as *z* ∈ ℝ^*N×K*^ where setting *z* = *A* at the reparameterization step guarantees an archetypal latent space. In this, *z*_*i*_ = *A*_*i*_ where *A*_*i*_ ∈ Δ_*K*−1_, which states that cell weights must be archetypal via valid barycentric coordinates on the Δ_*K*−1_ simplex such that

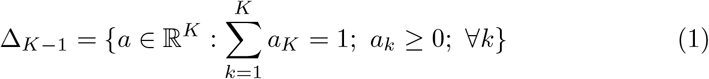

These constrains are enforced at the reparameterization step during training, updating

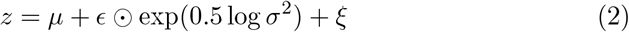

where *ϵ* ~ 𝒩 (0, *I*) and *ξ* ~ 𝒩 (0, 0.05^2^*I*) represent standard reparameterization noise and an added exploration noise term to prevent numeric collapse at boundary conditions, respectively. In this, *z* is subject to barycentric constraints via the user-selectable activation functions for either barycentric mode where *z* = softmax(*z*) or ReLU mode, with 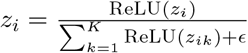 setting up row-wise normalization where *ϵ* = 10^−8^ prevents division by zero. These constraints follow the original formulation of archetype analysis, such that

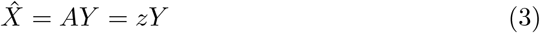

where each reconstructed sample is a convex combination of archetypes 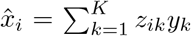. During training, *z*’s values are updated through archetypal reconstruction, implemented as described in [Gallo et al],

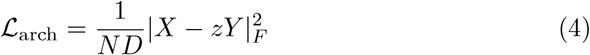

to minimize reconstruction error across all convex combinations of archetypes simultaneously via the Frobenius norm. This implementation was chosen to ensure smooth differentiable gradients across all data regardless of input matrix dimensionality or sparseness. The Frobenius norm squared,

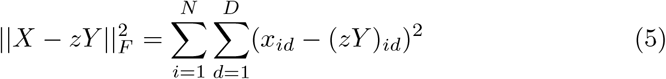

where *x*_*id*_ is the value of feature *d* in sample *i* and 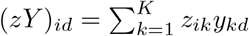 is the reconstructed value, is equivalent to the sum of squared reconstruction errors across all cells and features. This then provides smooth gradients in the form of

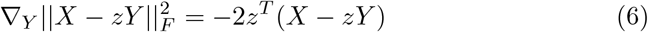

where *z*^*T*^ ∈ ℝ^*K×N*^ is the transposed coordinate matrix and the result is a *K × D* gradient matrix. Archetypal loss, constraint violation, and archetype stability are tracked across training epochs. Constraint violations are calculated every *M* epochs to track sum-to-one errors and non-negativity in the *A* matrix with a default *τ* = 10^−3^ tolerance threshold while archetype stability is tracked per-epoch on a per-archetype basis as 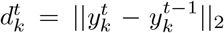 as well as across archetypes for mean drift as 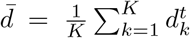 and maximum drift 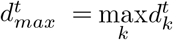. Over a 5-epoch window, position variance is also tracked as 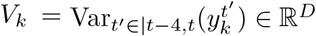 based on per-archetype variance 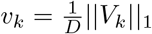 and mean variance 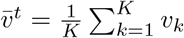. These terms measure how much archetype positions projected to PCA space shift during training. These tracking metrics help measure convergence, where low drift 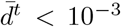 indicates convergence, low variance 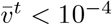 indicates stable archetype positioning, and high maximum drift 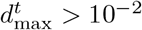 suggests archetype position instability and non-convergence. Parameters are optimized via Adam with a default learning rate of *α* = 0.001 and batch size *b* = 128 with early stopping monitoring archetypal on validation data where default patience is set to 10 epochs and training improvement threshold is set to 10^−4^ to invoke ReduceLROnPlateau scheduling. Training is deterministic (seed = 42) for reproducibility and includes updates to the input AnnData object for training history and other attributes.

Model quality is assessed via variance explained by implementing archetypal loss-based reconstruction, henceforth termed archetypal *R*^2^, as

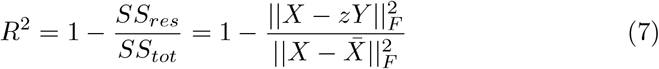

where *SS*_*res*_ is the residual sum of squares, *SS*_*tot*_ is the total sum of squares, and where 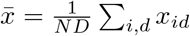 is the global mean, and 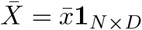 is that scalar mean value broadcast across all entries. The Deep_AA model provides an initialize_with_pcha option, which implements the Mørup and Hansen (2012) principal convex hull analysis algorithm 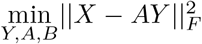 to quickly find the corners of the data, subject to 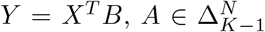, and 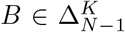 for K archetypes where A represents the data’s archetypal coordinates and B represents the archetype’s data coefficients. This can be used to initialize archetypal positions at the edges of the data manifold. Combined with PCHA initialization, a scalar inflation parameter can be set to evenly inflate all archetype positions away from the global centroid. This is implemented as

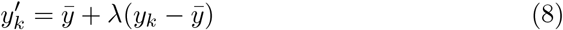

where 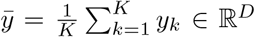 is the archetype centroid, *λ >* 1 is the inflation factor, and 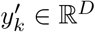 is the inflated archetype position. If PCHA fails, archetypes are initialized using a greedy furthest-sum strategy based on the initial step of PCHA. Another option for use_hidden_dims is also provided, which implements a nonlinear transformation of the archetypes before reconstruction

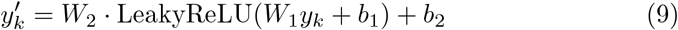

for each archetype *k* where *W*_1_, *W*_2_ ∈ ℝ^*D×D*^ are learnable transformation matrices and *b*_1_, *b*_2_ ∈ ℝ^*D*^ are bias vectors. This parameter is also exposed in the hyperparameter grid search function, which is designed to identify the optimal K archetypes that fit the data best, via the CVTrainingManager class. Three hyperparameters are exposed for exploration: archetype number *K*, encoder architecture depth (hidden layer dimensions), and archetype inflation factor *λ*. The search space is configured via a SearchConfig object specifying ranges for each hyperparameter. Cross-validation is implemented with k-fold splitting (default *k* = 3) to ensure robust performance metrics across data partitions. Each configuration is trained on (*k* − 1)*/k* of the data and validated on the held-out fold, with performance metrics aggregated across all folds. To manage computational costs on hyperparameter search in large datasets, PEACH implements a subsampling strategy that samples a maximum of 15,000 cells per fold using stratified selection based on categorical annotations in the AnnData .obs category to ensure proportional representation of cell types or conditions in each fold. Training with cross-validation folds employs early stopping with configurable patience (default 5 epochs), which monitors validation archetypal *R*^2^ with a minimum improvement threshold of 10^−4^. Training epochs for cross validation are typically restricted (default max_epochs_cv = 15-25) compared to final model training of 50-100 epochs. For each configuration, grid search tracks archetypal *R*^2^ as the primary metric while also tracking root mean squared error (RMSE), mean absolute error (MAE), convergence epoch count, and cross-validation stability as measured via the standard deviation of archetypal *R*^2^ across folds. Results are compiled into a CVSummary object supporting three core decision support methods: 1) rank_by_metric() for flexible ranking of any specified tracked metric (this enables nomination of top ranked hyperparameters via code in script-based analyses or pipelines), 2) summary_report() for consolidated performance summaries, and 3) visualization options plot_elbow_r2() and plot_metric() to visualize archetypal *R*^2^ or any other tracked metric, respectively. PEACH deliberately separates the hyperparameter search step from the final model training step to provide users with robust options to choose the configuration best suited to their data and use case.

Additional optional loss metrics are provided, including a standard KL divergence term, a diversity loss term to encourage separation between arguments, a regularity loss term to encourage balanced usage of all archetypes, a sparsity loss term to encourage sparse archetypal representations, a manifold regularization loss term combining a nearest neighbor penalty, a bounding box penalty, and a density penalty to encourage the archetypal representation to stay near the data manifold. The total loss term can be expressed as

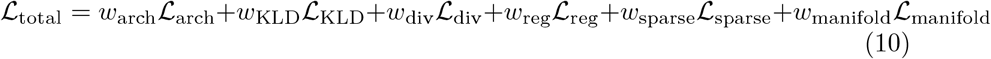

In this, the KL divergence term is implemented as

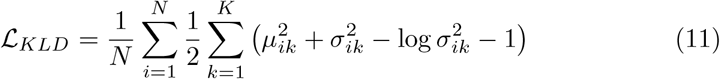

where *µ*_*ik*_ is the mean of archetype coordinate *k* for sample *i* and 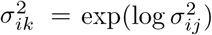 is the variance. Diversity loss is implemented as

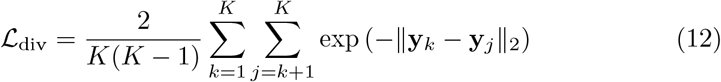

where **y**_*k*_, **y**_*j*_ ∈ ℝ^*D*^ are archetype positions, ∥ *·* ∥_2_ is the Euclidean norm, and 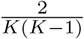 normalizes over all unique archetype pairs, which penalizes pairs of archetypes that are too similar. Regularity loss is implemented as

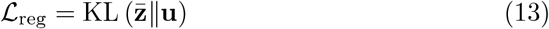

where 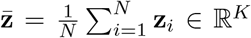 is the mean archetype usage across the dataset, 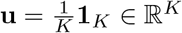 is the uniform distribution over archetypes, and 1_*K*_ is a vector of ones, which prevents the model from ignoring some archetypes. Sparsity loss is implemented as

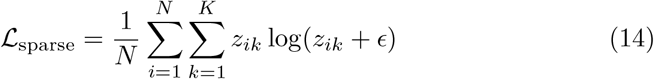

where *z*_*ik*_ are the archetypal coordinates, *ϵ* = 10^−8^ prevents log(0), and the entire expression is the negative entropy to penalize cells that spread across many archetypes. Manifold loss combines 3 components, a) the nearest neighbor penalty expressed as

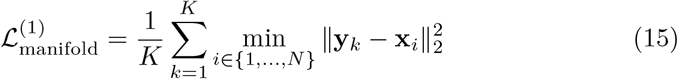

where **y**_*k*_ ∈ ℝ^*D*^ is the *k*-th archetype, **x**_*i*_ ∈ ℝ^*D*^ is the *i*-th data point, and the min operation finds the closest data point to each archetype, b) the bounding box penalty is implemented as

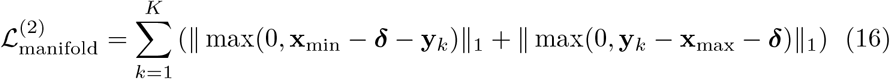

where **x**_min_, **x**_max_ ∈ ℝ^*D*^ are element-wise minimum and maximum values across the dataset, ***δ*** = 0.1(**x**_max_ **x**_min_) ∈ ℝ^*D*^ is a margin vector, max(0, ·) is elementwise, and _1_ is the L1 norm to ensure archetypes stay near the data’s bounding box range, and c) a density penalty implemented as

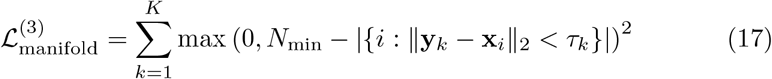

where *τ*_*k*_ = 1.5 median_*i*_( ∥ **y**_*k*_ − **x**_*i*_ ∥ _2_) is an adaptive distance threshold for archetype *k* using |{·}|as set cardinality (number of nearby points) and *N*_min_ = max(1, 0.05*N*) sets the minimum required nearby points to penalize archetypes that are far away from most data. Manifold loss terms are combined as 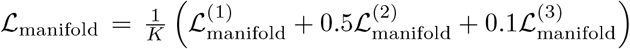. All loss terms beside archetypal loss are set to 0 by default but available for exploration if the data set being trained on calls for alternative feedback mechanisms. Archetypal loss *R*^2^ of 0.9 and KLD loss of 0.1 has provided high performance for many real scRNAseq datasets explored thus far.

Also included are additional training metrics, including a standard root mean squared error implemented

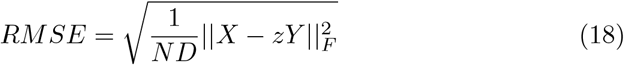

where *N* is the number of samples, *D* is the number of features, and normalization by *ND* provides interpretable reconstruction error in the original data scale. Archetype usage is assessed via archetype entropy, implemented as

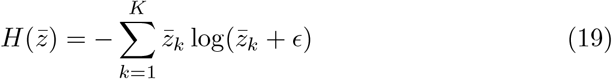

and sparsity, implemented as

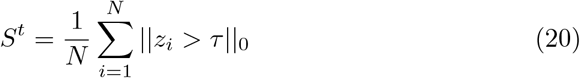

with *τ* = 0.01 default, where 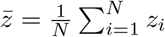 is mean archetype usage.

### Accessing archetypal attributes

Archetype analysis operates in PCA space and the relationships between cell and archetype coordinate positions can be used for develop statistical interpretations of biological characteristics associated with archetypal specialization or tradeoff. Calling .tl.train_archetypal() implements a full training loop implementing all of the above features and returns an AnnData with the trained model and the archetypal coordinates in PCA space stored in .uns. Training results are stored in a returned dict and available at the key ‘history’. Invoking .tl.archetypal_coordinates() then calculates pairwise distances in PCA space between all cells and all archetypes, stored in .obsm. Previous work in archetype analysis with biological data defines the 10-15% of cells closest to each archetype as in that archetype, which is implemented as .tl.assign_archetypes with the parameter precentage_per_archetype available to set the desired percentage of cells to include in each archetype group. This updates cell IDs with archetype identifiers in .obsm, with a global centroid group sampled for the same percentage of nearby cells as archetypes labeled as ‘archetype_0’ and all remaining cells labeled ‘no_archetype’. The global centroid is computed across all cells in the sample and typically represents the population of cells with the least degree of archetypal specialization. The above attribute access and update workflow sets up all archetype-cell distances and labels for further analysis in gene, gene set, and conditional variable association tests.

### Archetype gene enrichment

After binning cells by archetype, archetype-gene associations can be tested using a one-versus-all Mann-Whitney U test framework with multiple testing correction. For each archetype tested, cells are partitioned into 2 groups: those assigned to the archetype versus all other cells or all other archetypes, as selected by a user, to set up high and low groups for the non-parametric Mann-Whitney U test (also known as the Wilcoxon rank-sum test). The default test layer is log counts. The U statistic as calculated as 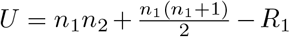 where *n*_1_ and *n*_2_ are the sample sizes of archetype and non-archetype cells respectively, and *R*_1_ is the sum of ranks for archetype cells after pooling and ranking all observations. The test is implemented with a two-sided alternative hypothesis (default) to detect both elevation and depletion, though users can specify one-sided tests when directional hypotheses are appropriate. Effect sizes are quantified through two complementary metrics: log_2_ fold-change computed as 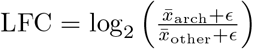 where *ϵ* = 10^−8^ prevents division by zero, and the rank-biserial correlation coefficient 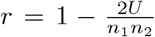 which ranges from −1 to 1 and measures the probability that a randomly selected archetype cell has higher expression than a randomly selected non-archetype cell. Multiple testing correction is implemented via Benjamini-Hochberg procedure to control the false discovery rate (FDR) at a user-specified threshold (default *α* = 0.05). PEACH provides 3 FDR correction scopes: ‘global’ correction across all tests, ‘per_archetype’ correction within each archetype separately, and ‘none’ for raw p-values. Global is recommended for initial analysis as it provides the most stringent correction across groups. Results are returned in a comprehensive DataFrame containing columns for feature name, archetype identifier, test statistic, raw p-value, FDR-corrected p-value, effect size (log fold change or mean difference), significance flag, and direction of change (higher/lower), which can be visualized in downstream dotplot functions for understanding archetype phenotypes.

### Gene set loading and AUCell activity scores

PEACH implements gene and gene set analysis via MSigDB gene set collections and single cell pathway activity scores from AUCell. Gene set networks are loaded via the load_pathway_networks() function, which supports all MSigDB collections and automatically handles organism-specific gene nomenclature for human and mouse datasets mapping gene symbols to match the input AnnData variable names. Pathway activity quantification via AUCell is implemented via compute_pathway_scores() which uses AUCell’s functions to rank genes by expression within each cell and calculate a recovery curve of each gene’s ranked positions, returning an area under the curve metric that is invariant to library size or gene count distributions^23^. Activity scores are stored in AnnData .obsm[‘pathway_scores’] to maintain consistent indexing. Statistical testing for archetype-gene set associations is then implemented exactly as described above for archetype gene enrichment using the Mann-Whitney U-test framework, where pathway result effect sizes are reported as the mean difference in scores as 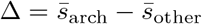 rather than log-fold change as the pathway scores are already on a normalized scale.

### Hypergeometric test

To identify enrichment of categorical variables, such as cell type or treatment stage, with archetypes, PEACH implements hypergeometric testing via a 2 *×* 2 contingency table framework. For each archetype-condition pair, cells are clarified by 2 binary variables: archetype membership (assigned vs not assigned) and condition status (condition present vs absent). The rest computes the probability of observing at least the observed number of in-archetype cells with the condition, given the marginal totals, under the null hypothesis of random assignment. This is calculated as

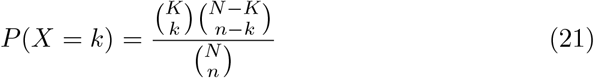

where *N* is the total number of cells, *K* is the number of cells in the tested condition, *n* is the number of cells in the archetype, and *k* is the observed overlap between archetype and condition. The p-value on this test is calculated as the right-tail cumulative probability 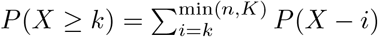 for each enrichment test. Effect size is quantified via the odds ratio

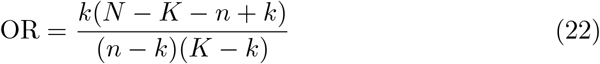

which measures the increase in odds of condition membership within the tested archetype compared to all non-archetype cells. An odds ratio of 1 indicates no association, OR > 1 shows enrichment, and OR < 1 shows depletion. Confidence intervals for the odds ratio are then computing using the Woolf method:

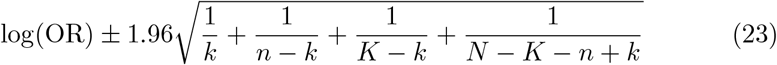

to provide statistical uncertainty estimates for the effect sizes. A continuity correction is added to handle edge cases where all cell counts in the contingency table cells approach 0 by adding 0.5 to all contingency table cells when any cell count is less than the minimal threshold (default 5 cells). Multiple testing correction follows the same Benjamini-Hochberg FDR framework as the gene association testing, with correction applied across all archetype-condition pairs. Results are returned via DataFrame containing archetype identifier, condition value, observed and expected cell counts, odds ratio, 95% confidence intervals, raw p-values, and FDR-corrected p-values.

### Advanced pattern analysis

The nature of a rotating 1-vs-all Mann-Whitney U-test means that the same gene or pathway can be associated with multiple archetypes, revealing potential grouped tradeoff patterns of interest across the archetypal space. To support this, PEACH implements three specialized pattern analysis functions to identify archetype-associated features with complex relationships across multiple archetypes. These are each designed to address a different biological question. First, in archetype_exclusive_patterns(), a more stringent test between each archetype *A* and each feature *f* evaluates whether *f* is significantly elevated in *A* compared to every other archetype individually. A feature is considered exclusive to the archetype *A* if and only if it satisfies 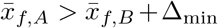 for all *B* ≠ *A* where 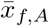 is the mean expression in archetype *A* and Δ_min_ is a minimum effect size threshold (default 0.05). In other words, is feature *f* high only in archetype *A*? Statistical significance is assessed via pairwise Mann-Whitney U tests with Benjamini-Hochberg FDR correction applied across all pairwise comparisons. This function returns pathways that are uniquely enriched in each archetype and not shared across other archetypes. Second, specialization patterns are identified via specialization_patterns(), which tests features against the global centroid archetype_0 to differentiate what distance-dependent features distinguish each archetype from the undifferentiated global center of the data. This tests 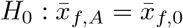 against 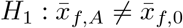 for each archetype and feature *f* using Mann-Whitney U tests between archetype-assigned cells and global centroid-assigned archetype_0 cells. Features with positive effect sizes indicate upregulation upon specialization towards the tested archetype while features with negative effect size indicate downregulation. Finally, third, mutual exclusivity patterns can be identified using tradeoff_patterns(), which can operate in two modes: pairwise tradeoffs for binary antagonism and complex patterns for multi-archetype tradeoff patterns. In pairwise mode, the function tests all unique archetype pairs (*A, B*) to identify features exhibiting antagonistic expression patterns: elevated in and suppressed in *B* or vice versa, to identify Pareto fronts of tradeoff features. For each pair and feature, a Mann-Whitney U test is used to compare the distribution of the feature in versus *B* directly with effect size quantifying the magnitude of the difference. Results include pattern codes in the format Axxxx_xxBxx indicating archetype *A* is high and archetype *B* is low for a 5 archetype system to summarize which features are associated with which archetype tradeoff patterns. In complex pattern analysis mode, the function generates all possible high/low archetype combinations up to a maximum pattern size (default 3 archetypes per group), testing whether features show consistent elevation in one archetype subset against consistent suppression in another, to identify regulatory features and tradeoff patterns that span larger regions of the archetype space. For example, a pattern 12x_xx3 indicates elevation in archetypes 1 and 2 with suppression in archetype 3. All pattern analysis functions return standardized DataFrames with consistent column names: feature name, archetype identifier(s), effect size, Mann-Whitney U test statistic, raw p-value, FDR-corrected p-value, significant flag, direction indicator, and pattern_code. Effect size thresholds (min_effect_size parameter) filter for biological relevance. FDR correction is applied as described above.

### Visualization

PEACH provides comprehensive visualization functions for all features described above. The primary visualization function pl.archetypal_space() generates interactive .html 3D scatterplots of cells in archetypal coordinate space using the first 3 PCA components, colored by user-specified variables, such as archetype assignment, other categorical variables in the AnnData .obs slot, or gene expression levels. A helper function to co-visualize multiple archetype fits is provided in visualize_archetypal_space_3d_multi() for comparing archetypal fits. Statistical test results are visualized through pl.dotplot(), which generates summaries for any of the feature tests described above using 4 dimensions of information: features (y-axis), archetypes or pattern_code (x-axis), effect size (dot size) and statistical significance (dot color). This function provides a top_n_per_group parameter to size each group. Hyperparameter search results can be visualized from 2 CVSummary methods, where pl.plot_elbow_r2 is dedicated to visualization of archetypal *R*^2^ across training epochs and can be faceted to visualize performance across several input hyperparameter ranges. The generic pl.plot_metric() function applies the same approach to any tracked metric (RMSE, MAE, convergence epochs, etc) with automatic legend generation. Training diagnostics can be visualized through pl.training_metrics(), which generates a multi-panel summary of loss components, performance metrics, and archetype stability metrics over training epochs that was included to track hyperparameter configuration results. Additional visualization functions support specialized analyses. 2D pairwise scatterplots of archetype coordinates across principal components are available via pl.archetype_positions() while pl.archetype_positions_3d() provide a matplotlib alternative view of archetype positions without a cell scatterplot overlaid. The dotplot framework is extended to advanced pattern analysis via pl.pattern_dotplot() and pl.pattern_heatmap() with enhanced clustering and filtering options.

## Results

### Pooled myeloid cells hyperparameter search

To test PEACH, we pulled 263,159 human scRNAseq profiles from normal hematopoiesis in circulating blood^22^. Using author-provided annotations, we split this dataset down to only myeloid cell types, covering common myeloid progenitors, dendritic cells, granulocyte monocyte progenitor cells, promonocytes, conventional dendritic cells, CD14+ monocytes, plasmacytoid dendritic cells, pre-conventional dendritic cells, and CD14+/CD16+ monocytes. This subset comprises 84,566 cells with 28,121 gene expression features. Data had been previously preprocessed for logcounts, normalization, and filtering of low quality reads and high mitochondrial cells. PCA was performed to identify the minimal number of PCs with higher than 95% variance explained: 13. Data was not filtered down to highly-variable genes only. Picking up at the PCA-projected data, we applied the hyperparameter grid search feature from PEACH to test 3:13 archetypes and hidden_dims options of [128, 256, 512], [128], [64, 128, 256], and [64, 128]. The grid search first subsamples datasets larger than 15,000 cells via random permutation, then performs 3-fold cross-validation on this data. For each fold, independent train (67%) and validation (33%) indices are created as separate TensorDatasets and DataLoaders. Each fold is trained for up to 50 epochs (balanced preset) with early stopping based on validation performance, using PCHA-based initialization of archetype positions. Archetypal *R*^2^, measured via a Frobenius norm implementation of data reconstruction from archetype weights, and root mean squared error (RMSE) were calculated for each configuration. After hyperparameter search, results were summarized in elbow plots for visual inspection and selection of the smallest k archetypes with high archetypal *R*^2^ (**Figure 1A**). From this, we selected 7 archetypes and hidden_dims [128, 256, 512] for final training. On 84,566 cells running on CPU, this hyperparameter search process required approximately 30 minutes to complete, representing a computationally tractable hyperparameter search function for large scale archetype analysis.

**Figure 1:**
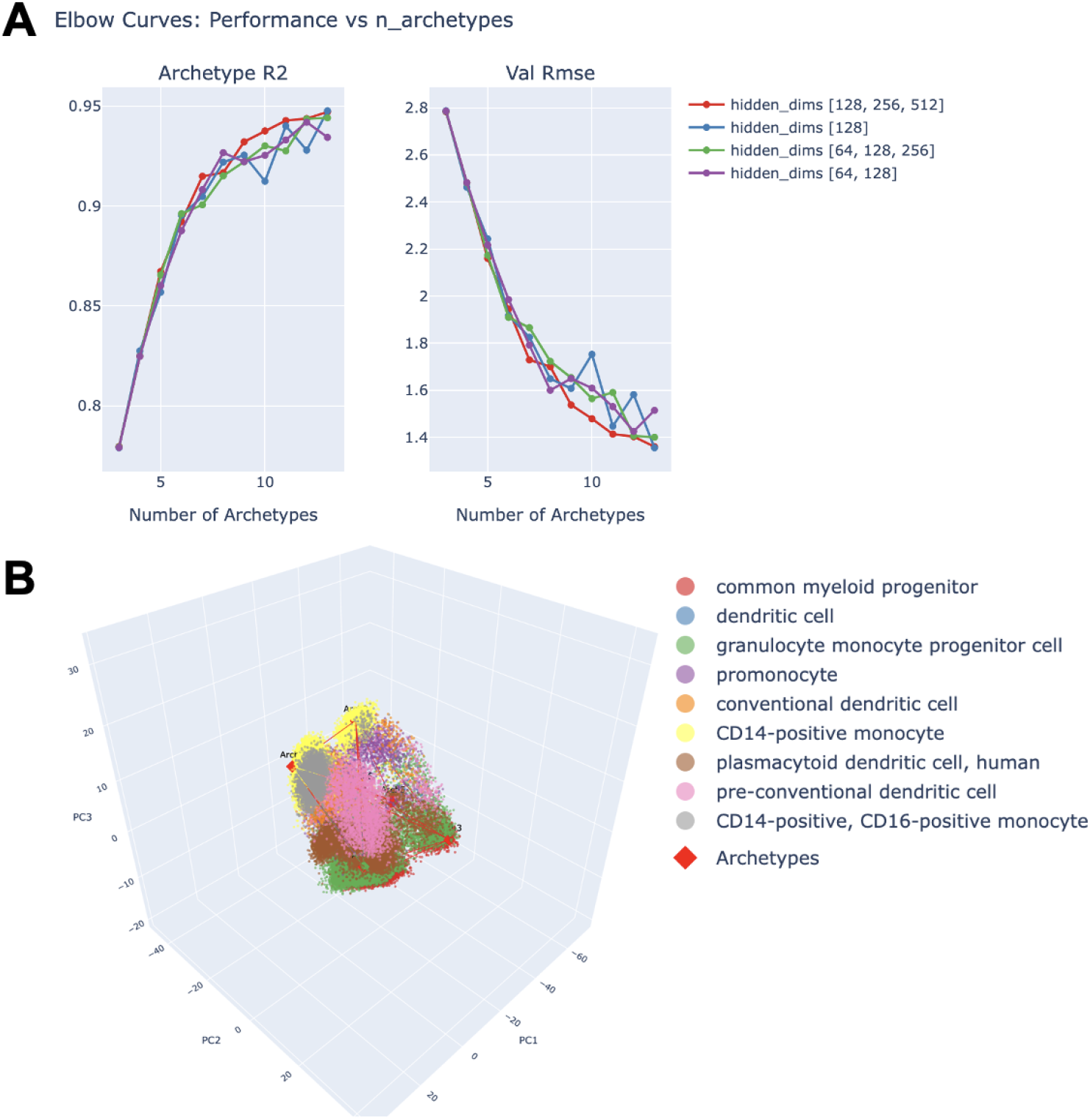
Hyperparameter search and 7 archetype fit in pooled myeloid cell types. **A:** PEACH hyperparameter search implements a 3-fold cross-validation framework for 25-100 epochs of PEACH-based archetypal analysis for each configuration specified. Shown are the resulting elbow curves for archetypal *R*^2^ and root mean squared error (RMSE) showing archetypal reconstruction and summary model error for 3 to 13 archetypes and a variety of hidden dimension configurations. From this, a configuration of 7 archetypes and [128, 256, 512] hidden dimensions were chosen. **B:** A final PEACH fit based on the settings from 1A, showing archetype coordinates as red dots, hull connections via red lines, and cell types each colored individually in PCA space. As expected with mixed cell types, archetype analysis recapitulates cell-type specific separation of phenotypes.

### Archetypes of pooled myeloid cells

Trained for 200 epochs, 7 archetypes on pooled myeloid cell type scRNAseq data resulted in a final archetypal *R*^2^ of 0.9343, showing that input data is well-reconstructed from learned archetypal weights, with a final root mean squared error (RMSE) of 1.5119. In this, archetypal loss weight was set to 0.9 and all other loss terms were set to 0. Exploration of other loss configurations has shown the greatest impact from a modest weight on the diversity loss term <0.1 The pooled myeloid cell types archetypal fit is shown in **Figure 1B**, where cells are colored by cell type, which shows that cell types largely separate out at archetypes, reflecting phenotypic differences in cell type emerging as archetypal specialization. Archetypal hulls may sometimes fit partially within the training data when both are projected to PCA space due to initialization conditions and the archetype-cell position feedback loop central to training, which updates archetypal positions via gradient descent on reconstruction loss. For these cases, an inflation_factor parameter is provided to uniformly increase archetype initialization distances from the global centroid of the initial archetype positions. Applying the built-in hypergeometric test, which estimates overrepresentation of a categorical variable in cells binned in a given archetype, to pooled myeloid cell types shows a strong association of each cell type with one or more archetypes (**Figure 2**). This function can be applied to attributes such as treatment status, timepoint, or sample source, but in this case underscores that archetype analysis is best-suited for analysis of phenotypic range and tradeoffs in a single cell type at a time. Note that ‘archetype_0’ is not a functional specialist but is instead binned from the 15% of cells closest to the global data centroid, representing the most highly-mixed, least-specialized cell state present in the sample. This is calculated to enable trajectory-based analyses of specialization paths from the center of the data to each archetype.

**Figure 2:**
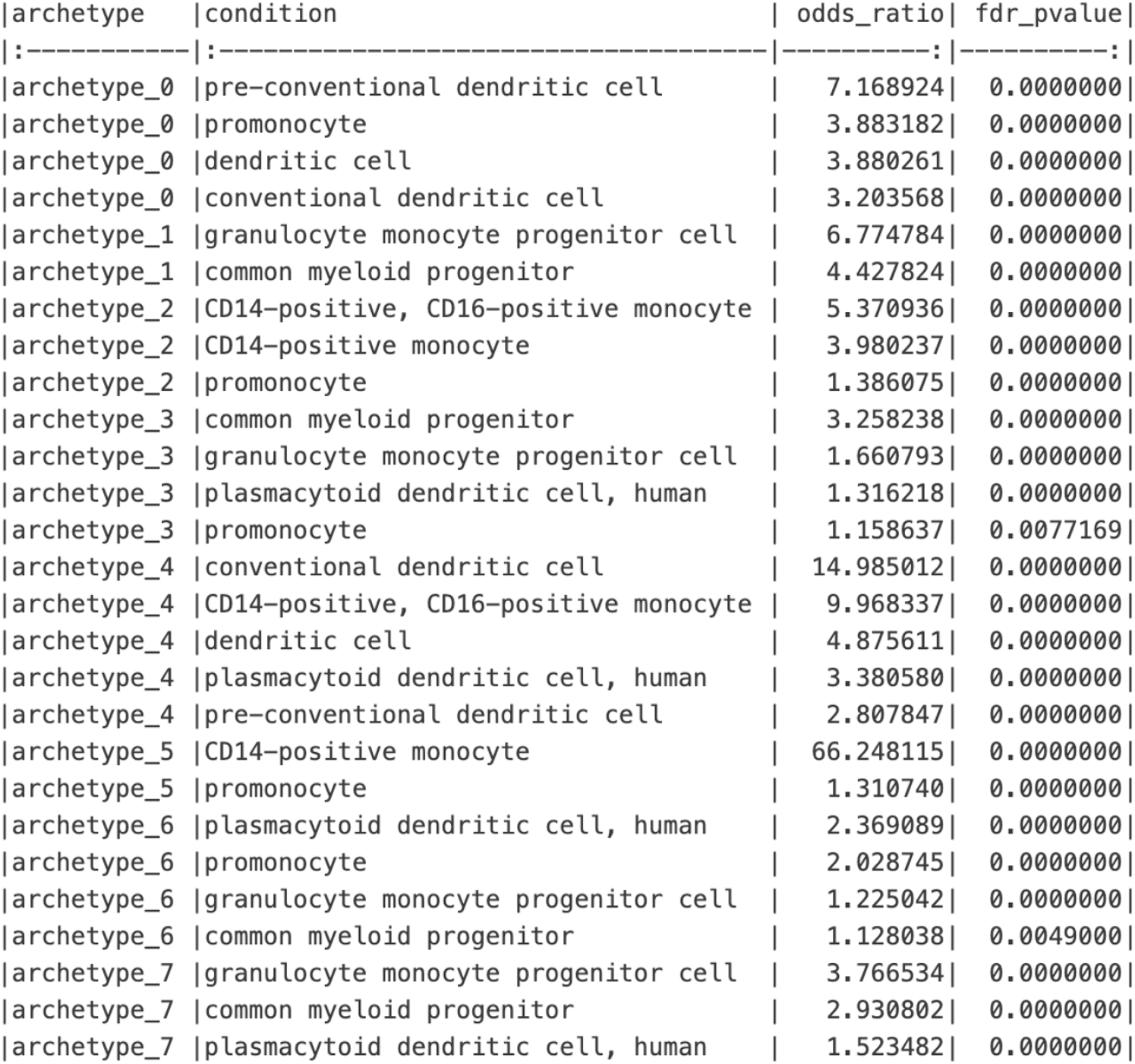
Hypergeometric testing in pooled myeloid cells reveals cell type-specific archetype associations. Running a hypergeometric test for dataset-derived cell type definitions against archetype bins (15% of cells closest to each archetype in PCA space) shows that, for example, granulocyte monocyte progenitor cells and common myeloid progenitor cells are significantly associated with archetype 1 at odds ratios for enrichment at 6.77 and 4.42, respectively. Archetype_0 is sampled from the 15% of cells closest to the global centroid, representing cells not associated with any archetype. Running archetype analysis on mixed cell populations often recapitulates clustering results with separation of distinct cell types at each archetype, although some cell types, such as promonocytes, may be associated with multiple archetypes and represent a cell phenotype subspace that can be more accurately characterized in archetype analysis run on a cell type-specific subset of the data.

### Plasmacytoid dendritic cells (pDCs) archetypes

Next we subset the pooled myeloid cell types down to a single cell type, plasmacytoid dendritic cells (pDCs) for a more focused investigation. In 9,328 pDC scRNAseq profiles, we re-ran the hyperparameter grid search to identify the best number of archetypes with high explanatory power as the best number of archetypes in a subset of data may not match that in the superset it is drawn from. This resulted in a configuration of 6 archetypes, hidden dims [64,128], and an inflation factor of 1.0, producing an archetypal *R*^2^ of 0.9488 and RMSE of 1.3458 over 200 training epochs (**Figure 3A**). From this, archetype coordinates, archetype weights, and cell-archetype distance attributes were calculated and added to the AnnData. Gene set scores were then calculated for every cell using the C5:BP set of gene set definitions and stored in the AnnData as well. Archetypes were defined as the 0.15 proportion of cells closest to each archetype and labeled in .obs, enabling comparison of genes and gene sets with each archetype using the Wilcoxon rank sum test (**Figure 3B**).

**Figure 3:**
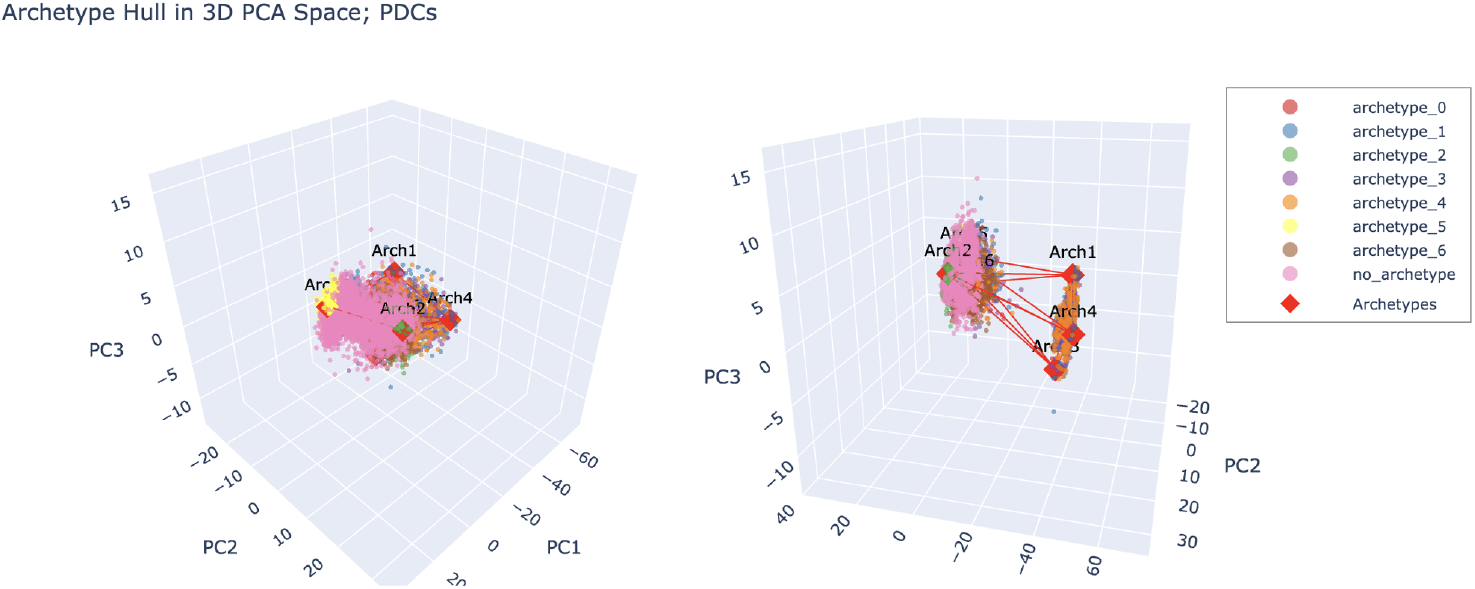
Plasmacytoid dendritic cell phenotypic diversity spans six archetypes separated by activation state. A subset of pure plasmacytoid dendritic cells (pDCs) can be reconstructed from weights for 6 archetypes, showing functional separation by cell activation markers for active proliferation and translation. This separation is apparent through all myeloid cells in the dataset, where each activation state is captured by 3-4 specialist archetype phenotypes representing metabolic adaptations, immune interactions, or active chemotaxis.

These results showed strong ribosomal protein gene enrichment across half of all archetypes, which were also associated with markers of ribosomal biogenesis, ATP synthesis, and active proliferation, in contrast with the other half of archetypes, which exhibited more immunoregulatory and quiescence-associated pathways. Notably, this contrast was present across all myeloid cell types in the hematopoietic dataset, reflecting a potential difference in cell activation status visible in PCA projection as well as archetype-associated functional attributes like genes and gene sets (**Figure 4ABC**) with cells projecting above PC1 −20 showing significant enrichment of ribosomal biogenesis and other activation markers. In pDCs, archetypes 1, 3, and 4 are associated with a quiescent state while archetypes 2, 5, and 6 are associated with an activated state featuring high ribosomal biogenesis (**Figure S1**). From pathway analysis and Wilcoxon rank sum testing, archetype 1 is associated with cap hypermethylation and ATP synthesis all driven by metabolic markers such as ATP5MC2/MG, H2AFZ, TUBA1B, and PCLAF, suggestive of a cycling cell state. Archetype 2 maintains active biosynthesis signatures with pathways in intrinsic apoptosis regulation, ribosomal biogenesis, and protein turnover with heavy enrichment of RPS/RPL genes, suggesting an activated, highly proliferative phenotype. Archetype 3 is associated with viral latency and cytoskeletal remodeling driven by TUBB, STMN1, ACTG1, and GZMB, associated with suppression of T-cell expansion^24^. Archetype 4, in contrast, is associated with regulation of p53-mediated apoptosis and protein degradation regulation, suggesting active translation. Archetype 5 shows enrichment for regulation of TLR7 and type 1 hypersensitivity with chromatin markers H3F3A and HIST1H4C upregulated, suggesting epigenetic modulation of inflammatory responses^25,26^. Finally, archetype 6 features the highest expression of ribosomal genes with active miRNA degradation, positive regulation of respiratory burst, and negative regulation of protein catabolism, suggesting an activated phenotype engaged with stressors and mounting active immune responses. These functional annotations are based on log-transformed gene count and AUCell-based gene set activity scores in a 1-vs-all Wilcoxon rank sum test, which treats each archetype as the high group and all other cells as the low group independently. This approach can lead to overlapping feature annotations at each archetype, requiring careful user discernment between results or the use of curated gene set subsets to investigate functions of interest.

**Figure 4:**
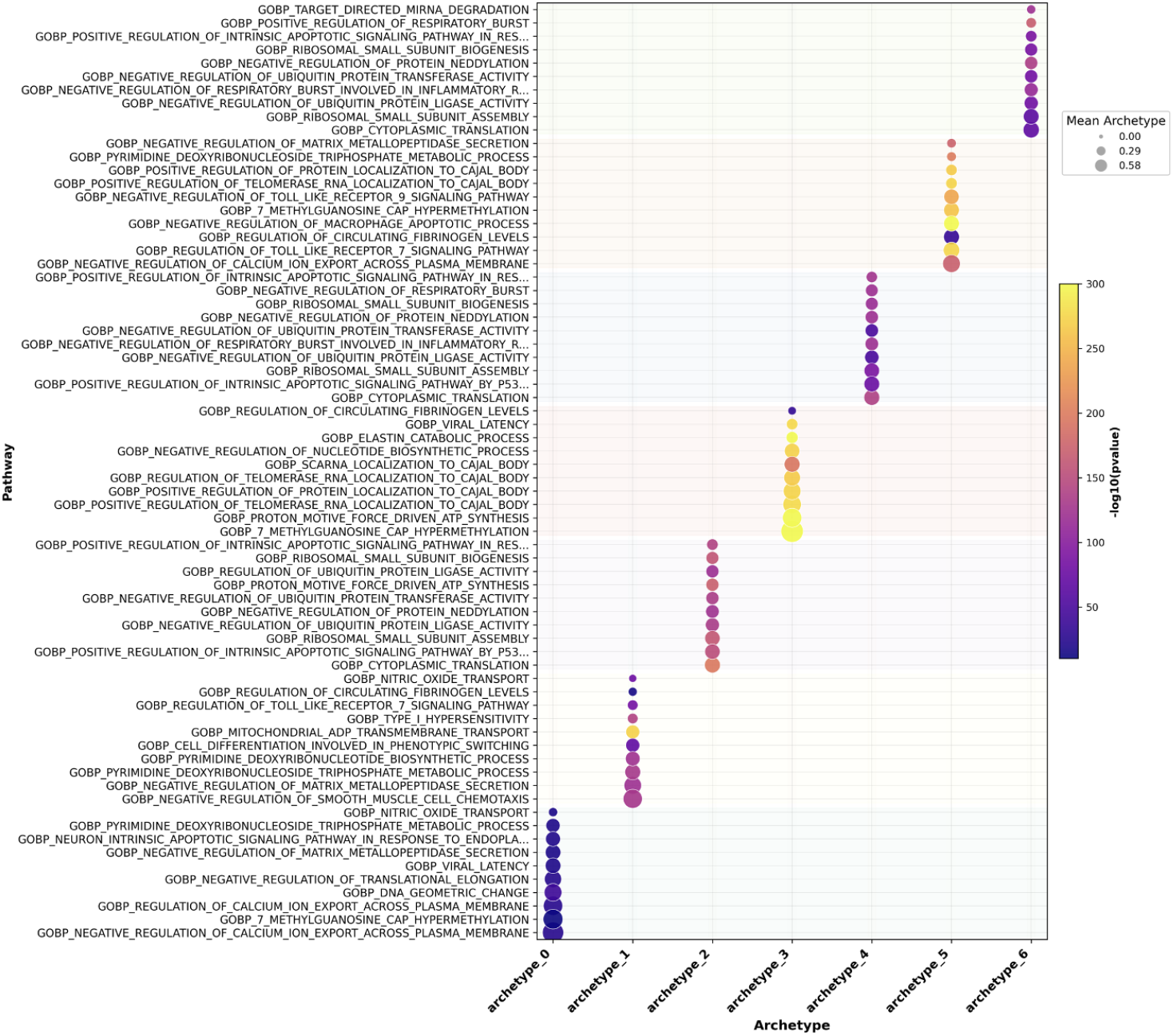
Plasmacytoid dendritic cells (pDCs) exhibit archetype-dependent functional specialization. Binning pDCs by the 15% of cells closest to each archetype and using a 1-vs-all Wilcoxon rank sum test for non-parametric association of archetype identity and gene expression or AUCell-based gene set activity scores reveals 6 archetype phenotypes spanning pDC diversity in this dataset.

### High-grade serous ovarian cancer treatment response trajectories separate by chemotherapy response score in archetype space

Next, we applied PEACH to the characterization of treatment response in longitudinal data using 12,097 epithelial ovarian cancer (EOC) cells subset out of non-spatial scRNAseq profiles from the *2024 Färkkila* study of immune cell changes in treated high-grade serous ovarian cancer (HGSOC)^27^. In brief, processed scRNAseq count matrices and metadata were obtained from the Gene Expression Omnibus, filtered for high mitochondrial and ribosomal content and low quality reads in Scanpy, and reannotated for cell types using PAX8, EPCAM, and WFDC2 as EOC markers in CellAssign. We ran Principal Component Analysis (PCA) on the normalized data for 50 components and selected the elbow curve at 11 PCs with >95% variance explained to use the least number of PCs with the highest explanatory power for archetype analysis, as it generally performs better without superfluous PCs with marginal variance explained. A hyperparameter grid search was performed as above, finding that 5 archetypes with hidden_dims [64, 128, 256] and an inflation factor of 1.75 resulted in an archetypal *R*^2^ of 0.9542 and RMSE of 1.1742 with early stopping triggered by convergence condition at epoch 191 (**Figure 5 inset**). Gene and gene set associations were tested as above. Gene set associations reveal an EOC archetype 1 specialized in antimicrobial immunity processes while EOC archetype 2 is specialized in immune engagement and antigen processing (**Figure S2**). EOC archetype 3 is specialized in protein deubiquitinylation and localization with suppression of natural killer cell cytotoxicity and EOC archetype 4 is associated with RNA processing and repression of translation elongation. EOC archetype 5, in contrast, is specialized in translation initiation and regulation of mitochondrial oxygen and electron transport. These associations are stable across multiple archetype analysis model fits, demonstrating strong reproducibility across model initialization conditions. Performing these association tests with gene set lists subset to functions of interest may reveal more module-specific archetypal specialization across the EOC phenotypic range. Notably, this dataset includes both pre-neoadjuvant chemotherapy samples as well as longitudinal post-chemotherapy (3-4 cycles of taxane and platinum chemotherapies) with chemotherapy response scores of ‘short’ for relapse and secondary surgical resection within 12 months of last platinum dose and ‘long’ for recurrence intervals longer than 12 months. From these annotations, we calculated centroid positions in PCA space for both long and short responders at each sample timepoint, showing that pre- and post-treatment trajectories diverge by chemotherapy response score in archetype space (**Figure 5**). Long responder pre-treatment centroids begin nearer EOC archetype 4 (RNA processing and translation elongation repression) and move towards archetypes 1 and 2 (immune engaged)–note that only disease recurrence samples are included here, indicating HGSOC that has evolved since last treatment where the pre-treatment phenotype may have been eliminated by treatment, leaving only minimally residual disease to recur after a phenotype shift. Short responder pre-treatment centroids begin at the point where long responders show recurrence, potentially demonstrating a more advanced or inflamed disease state less sensitive to treatment. Short responder trajectories then move further towards archetype 2, engaged in antigen presentation and immune cell interaction, including both activation and suppression of leukocyte cytotoxicity. Finally, we passed these conditions to the CellRank ConnectivityKernel by sampling the 15% of cells closest to each trajectory endpoint and characterizing the short and long responder trajectories individually, revealing that long responder trajectories are significantly associated, via the CellRank driver gene function, with S100A1/3/4/6, KRT19, UCA1, FTH1, SLPI, and ANXA2 whereas short responder trajectories are significantly associated with HLA-A/HLA-C/HLA-E/HLA-DPA1/HLA-DRB1, PPP1R15A, MT2A, BTG1, and SAT1, reflecting the short responder trajectory enrichment in the archetype 2 specialist phenotype. Interestingly, the long responder trajectory drivers, which are associated with later disease recurrence, reflect associations with the epithelial-to-mesenchymal transition and metastasis (S100A1/3/4/6), stress resistance (UCA1, FTH1, and SLPI), and angiogenesis (ANXA2).

**Figure 5:**
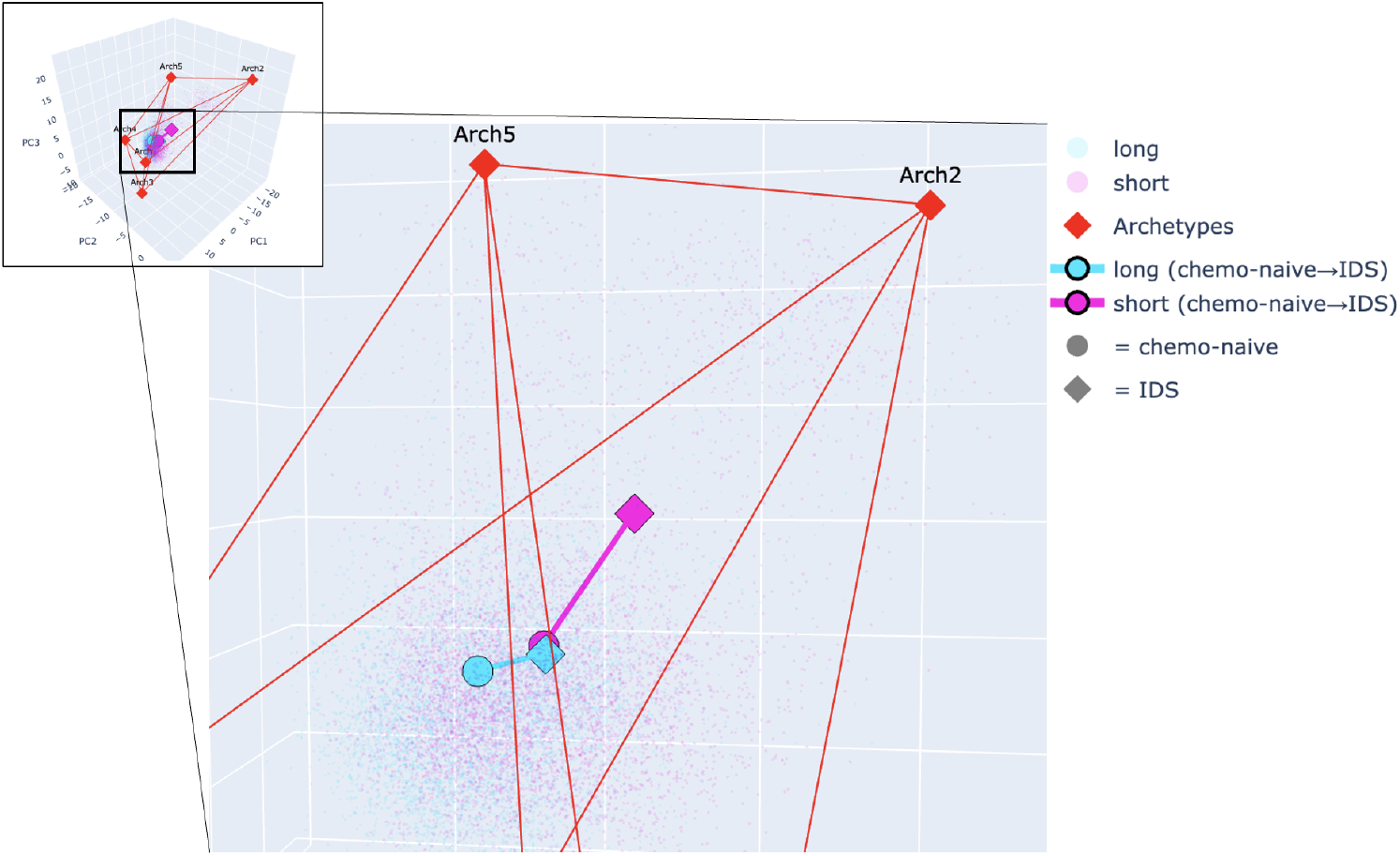
High-grade serous ovarian cancer cells follow different trajectories in archetype space. Inset: The global archetype analysis shows 5 archetypes spanning all high grade serous ovarian cancer scRNAseq profiles, reflecting metabolic and cell interaction tradeoffs in several pathways. Full: Zooming in on the divergence in pre-vs post-treatment trajectories by chemotherapy response score (CRS) shows that long responders begin at a distinct location, which corresponds to a particular set of specialist tradeoffs, and transition to the starting point of the short responders, potentially reflecting convergent evolution in treatment resistant cell states. Note that centroids across 22 patients are shown and represent the average behavior across myriad individual treatment trajectories.

## Discussion

Archetype analysis flips the assumptions of common clustering approaches to find the limits of phenotypic heterogeneity present in a sample and the tradeoffs between them. Instead of forcing every cell or sample into a discrete group with a clear border, archetype analysis instead identifies theoretical extremal states that bound the mixtures of specialist states present in real samples. This casts sample characterization in terms of cell state continua and varying mixtures of specialization weights, where distances from each archetype or direction between them carry interpretable biological information. Python Encoders for Archetypal Convex Hulls (PEACH) provides a PyTorch-based implementation of this framework that is compatible with the scVerse ecosystem, scalable to large modern single-cell datasets, and accessible to users without extensive hyperparameter tuning expertise.

The core contribution of PEACH is the integration of archetype analysis into a deliberately constrained autoencoder architecture where the latent space dimensions directly represent archetypal dimensions. This design choice eliminates the need for both alternating step optimization towards an archetypal fit and a separate archetypal decomposition step while ensuring that the learned representation maintains a clear, globally convex simplex structure in which every cell or sample can be located by barycentric coordinates. This constraint prioritizes interpretable archetypal coordinates over the generative sampling capabilities of a full variational autoencoder with a Gaussian latent space, preserving the core functions of archetype analysis where biological features can be related to archetypal extremal states in terms of distance, mixture, or shift through conditions. The Frobenius norm-based reconstruction loss provides smooth, differentiable gradients compatible with PyTorch-based optimizer setup, addressing the *O*(*n*^2^) scaling limitations of classical alternating least squares implementations that previously restricted archetype analysis to smaller datasets. A key feature distinguishing PEACH from previous implementations is the hyperparameter search with cross validation, which can also be used in an automated fashion to select the best hyperparameters for full training. Selecting the appropriate number of archetypes to fit data has been a persistent challenge in archetype analysis, typically requiring manual comparison of different *k* archetypes and the results from each. PEACH’s CVTrainingManager systematically explores archetype counts, hidden layer architectures, and inflation factor hyperparameters while providing interpretable summary statistics and visualizations to guide final configuration selection. The stratified subsampling strategy in CVTrainingManager enables tractable hyperparameter search even on datasets of hundreds of thousands of cells, as demonstrated in the myeloid cell analysis where a hyperparameter grid search across 44 configurations completed in approximately 30 minutes on CPU. This automation and integration with the scVerse ecosystem with a clear tool API significantly lowers the barrier to using archetype analysis for single-cell and other datasets and reflects design choices for future work exposing PEACH functions as tools for agent-based machine learning research in emerging single-cell atlases.

Application of PEACH to plasmacytoid dendritic cells (pDCs) revealed 6 archetypes spanning a split in activation state with half associated with ribosomal biogenesis and active proliferation and the other half with quiescent, immunoregulatory phenotypes. Finding this split across multiple archetypes reflects the nature of the underlying 1-vs-all Wilcoxon rank sum test, which independently tests each archetype as the high group against all other cells and can find the same gene or pathway significantly associated with each archetype. This, however, can also be used as an advantage to explore multi-archetype modules that exhibit tradeoffs across larger regions of the phenotypic space, such as the activation split in pDCs. Future work will include development of more specific statistical tests to identify archetype-exclusive phenotypes and separate related biological modules by archetype distance to provide a more granular view of co-regulated functions supporting each specialist phenotype near an archetype vs regulatory patterns and tradeoffs across the archetypal space. Testing longitudinal scRNAseq profiles with PEACH also reveals that different treatment conditions and responses are associated with distinct regions of archetype space, with chemotherapy response score-based centroids revealing that short and long responders begin and end at disparate points in specialization tradeoffs. The association of post-treatment long responders with the pre-treatment short responders may reflect the development of treatment-resistant states over time, with short responders already exhibiting a resistant state while long responders only converged on that state after treatment and >12 month before disease recurrence. The inclusion of a compatibility bridge to CellRank functions for trajectory characterization enables identification of driver genes associated with different archetypal or conditional source and target states, such as the resistance-associated genes driving the long responder trajectories and the immune evasion genes driving the short responder trajectories.

Future work will include extensions to other data types, different metric spaces, and more extensive perturbation modeling. By providing a performant, accessible, and extensible implementation of archetype analysis powered by PyTorch and compatible with existing single-cell analysis workflows, PEACH enables researchers to move beyond clustering-based analyses into identification of phenotypic limits, tradeoffs between them, and regulatory constraints governing cell fate choices after perturbation.

## CODE AVAILABILITY

PEACH is available as an open-source Python package at https://github.com/xhonkala/PEACH.

**Figure S1:**
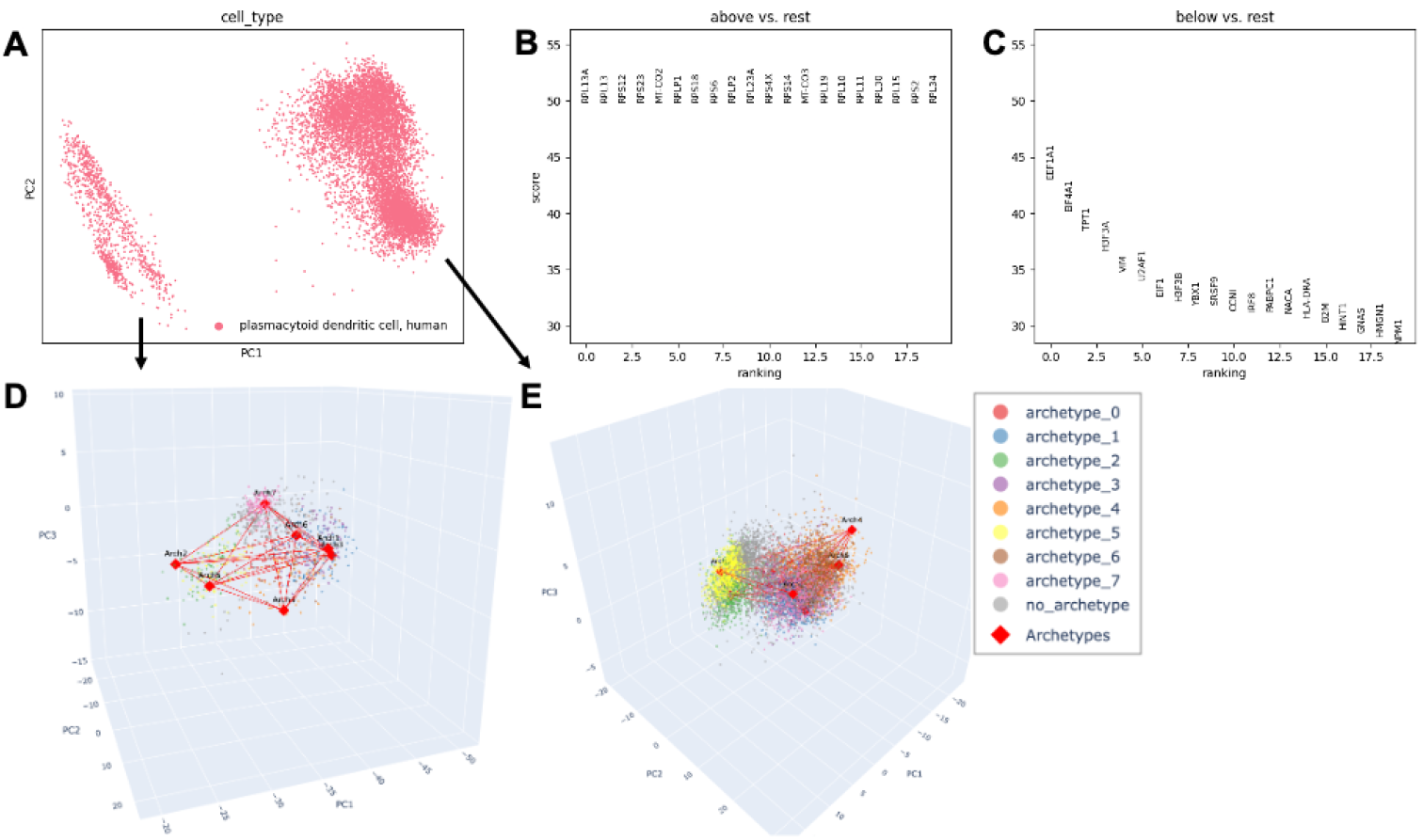
Plasmacytoid Dendritic Cells Exhibit an Activation-Based Phenotype Split. **A:** PCA projection of plasmacytoid dendritic cells (pDCs) showing a phenotypic split around PC1 −20. **BC:** Differential gene expression analysis from Scanpy (wilcoxon test) between high vs low groups, showing that the high group is dominated by ribosomal proteins reflective of active translation and proliferation while the low group is characterized by quiescence and immunoregulation. **D:** Low group independent archetype fit showing 6 archetypes across ~1500 cells. **E:** High group independent archetype fit showing 6 archetypes across ~6500 cells.

**Figure S2:**
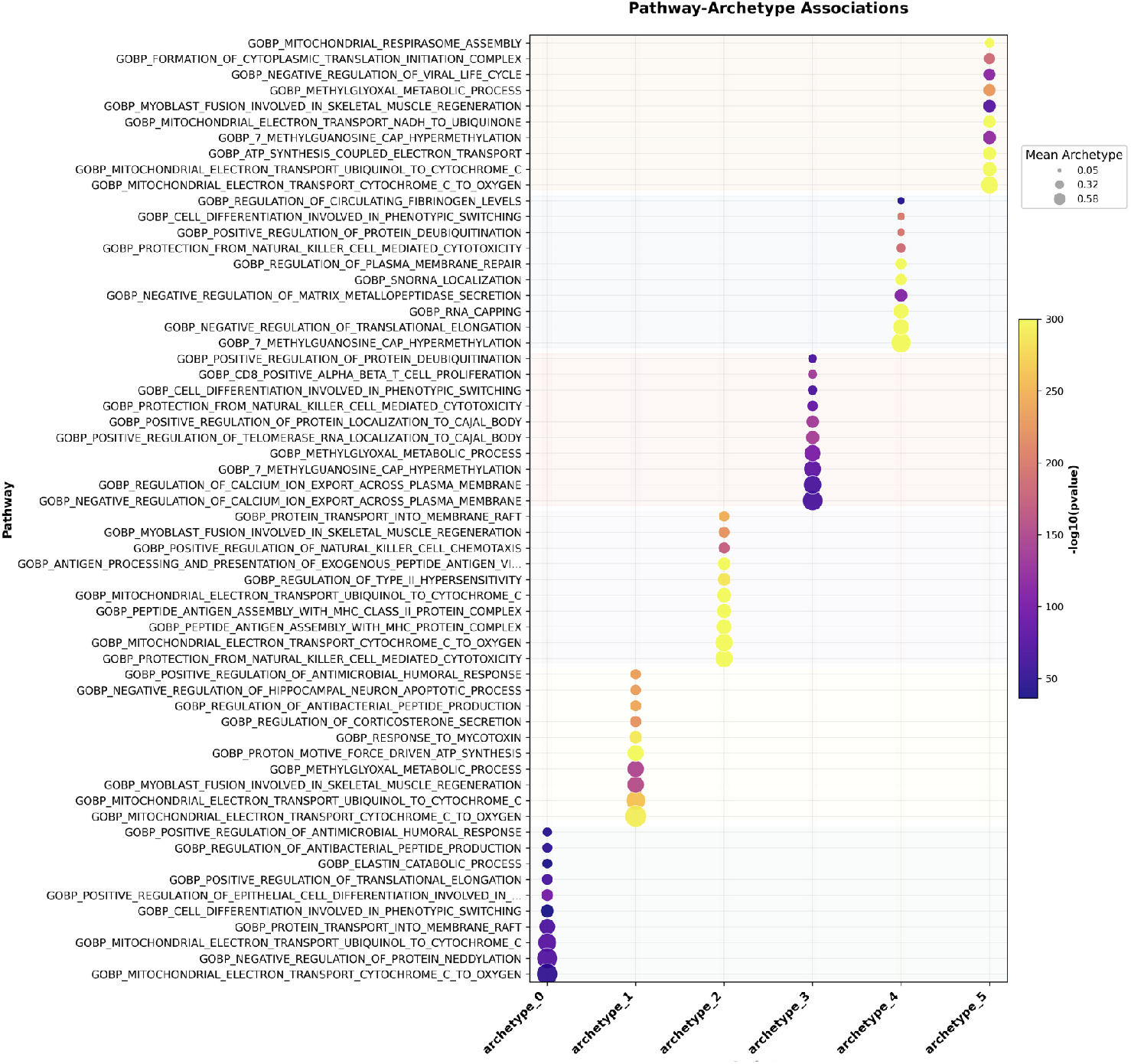
High-Grade Serous Ovarian Cancer Cell Specialist Phenotypes. The 15% of cells closest to each archetype were binned as in that archetype and set as the high group in independent 1-vs-all Wilcoxon rank sum tests for AUCell scores from C5:BP gene sets. This reveals different pathway associations across archetypes. Note that ‘archetype_0’ is a global centroid and is furthest away from all archetypes. Archetype 2 exhibits strong enrichment in immune engagement and regulation whereas archetype 4 is associated with RNA processing and translation regulation. More specific results can be obtained with function-specific subsets of gene sets tested. Future work will include the development of more specific archetypal phenotype statistical tests.

